# The H3.3K27M oncohistone antagonizes reprogramming in Drosophila

**DOI:** 10.1101/2020.11.10.375881

**Authors:** Kami Ahmad, Steven Henikoff

## Abstract

Development proceeds by the activation of genes by transcription factors and the inactivation of others by chromatin-mediated gene silencing. In some cases development can be reversed or redirected by mis-expression of master regulator transcription factors. This must involve the activation of previously silenced genes, and such developmental aberrations are thought to underlie a variety of cancers. Here, we express the wing-specific Vestigial master regulator to reprogram the developing eye, and test the role of silencing in reprogramming using an H3.3K27M oncohistone mutation that dominantly inhibits histone H3K27 trimethylation. We find that expression of the oncohistone blocks eye-to-wing reprogramming. CUT&Tag chromatin profiling of mutant tissues shows that H3K27me3 domains are globally reduced with oncohistone expression, suggesting that previous developmental programs must be silenced for effective transformation. Strikingly, mis-expressed Vg and H3.3K27M synergize to stimulate overgrowth of eye tissue, a phenotype that resembles that of mutations in Polycomb Repressive Complex 1 components. Our results imply that growth dysregulation can result from the simple combination of crippled silencing and transcription factor mis-expression, an effect that may explain the origins of oncohistone-bearing cancers.

## Introduction

Developmental programs in multicellular organisms are specified by transcription factors that activate and repress batteries of genes, thereby determining cell fate. Ectopic expression of specific transcription factors drive changes in cell fate, either by inducing pluripotency from a differentiated state [Takahashi & Yamanaka, 2006], or by transforming one cell type to another in a process referred to as ‘transdetermination’ or ‘direct reprogramming’ [Graf, 2011]. Aberrant reprogramming induced by transcription factor misexpression underlies some developmental and malignant diseases.

In eukaryotes transcription factors interact with chromatin, where genomic DNA is wrapped around histone octamers in nucleosomes. Silencing histone modifications on nucleosomes inhibit factor binding and transcription, and so modulate gene expression programs. Silencing has been suggested to impose directionality and reliability to developmental progression [Perino & Veenstra, 2016]. A key chromatin mechanism is mediated by trimethylation of the lysine-27 residue of histone H3 (H3K27me3) which is bound by Polycomb proteins. Mutations of Polycomb proteins derepress developmental transcription factor genes and thereby induce aberrant fate transformations in animals, plants, and fungi, highlighting the conserved importance of this chromatin system [Lanzuolo & Orlando, 2012; Bemer & Grossniklaus, 2012; Lewis, 2017].

In Drosophila, mutation of the histone H3-K27 residue recapitulates Polycomb transformations [Pengelly et al, 2013]. In humans, screening of cancer cells identified mutations of this residue in certain pediatric glioblastomas [Schwartzentruber et al, 2012; Wu et al, 2012]. These oncohistone mutations are lysine-to-methionine (K27M) mis-sense substitutions that dominantly inhibit the EZH1/2 histone methyltransferases and reduce chromatin methylation [Chan et al, 2013; Lewis et al, 2013; Bender et al, 2013; Justin et al, 2016; Stafford et al, 2018]. The oncohistone is not tumorigenic on its own, but may precondition cells to later oncogenic mutations [Pathania et al, 2017; Mohammad & Helin, 2017]. However, since the critical window for tumorigenesis is in early developing lineages, the sequence of initiating events is not accessible to analysis.

Histones and histone modifying enzymes are conserved across eukaryotes, and expression of the H3K27M oncohistone in Drosophila cells recapitulates chromatin and silencing defects seen in gliomas [Herz et al, 2014]. Here, we use Drosophila to show that the H3K27M oncohistone blocks direct reprogramming induced by ectopic expression of the wing master regulator transcriptional activator Vestigial (Vg). While oncohistone expression on its own inhibits cell proliferation, co-expression with Vg results in overgrowth of cells, and these cells retain eye identity. Chromatin profiling by CUT&Tag [Kaya-Okur et al, 2019] shows widespread reduction in H3K27me3 histone modification that may cripple silencing during reprogramming. These effects demonstrate that a defect in chromatin silencing combined with aberrant transcription factor expression is sufficient to induce neoplastic growth, with implications for the developmental origins of gliomas.

## Results

### The H3K27M oncohistone blocks direct reprogramming

To probe the interaction between reprogramming and chromatin silencing, we used inducible transgenes for the Vg master regulator transcription factor and for the H3.3K27M oncohistone. Vg encodes transcriptional activation domains and – when heterodimerized with the DNA-binding Scalloped (Sd) protein – determines the identity of cells in the pouch of wing imaginal discs [Ruiz-Losada et al, 2018]. Vg is a master regulator of wing development, as ectopic expression of Vg converts tissue into wing structures [Kim et al, 1996]. We used the *eyeless-GAL4* (*eyGAL*) driver to induce transgene expression during development of the eye [Hazelett et al, 1998]. Expression of Vg in the eye is semi-lethal (Table 1), and many dying pupae have small heads that lack eye tissue (Figure 1A,B). Those adults that survive show varying degrees of transformation: some animals lack eyes, while other have reduced numbers of eye ommatidia and have darkly pigmented outgrowths resembling wing tissue with small bristles and trichome-like hairs instead of the setae of normal eyes (Figure 1C; Table 1), indicating reprogramming of the eye by the Vg master regulator.

**Table 1.**
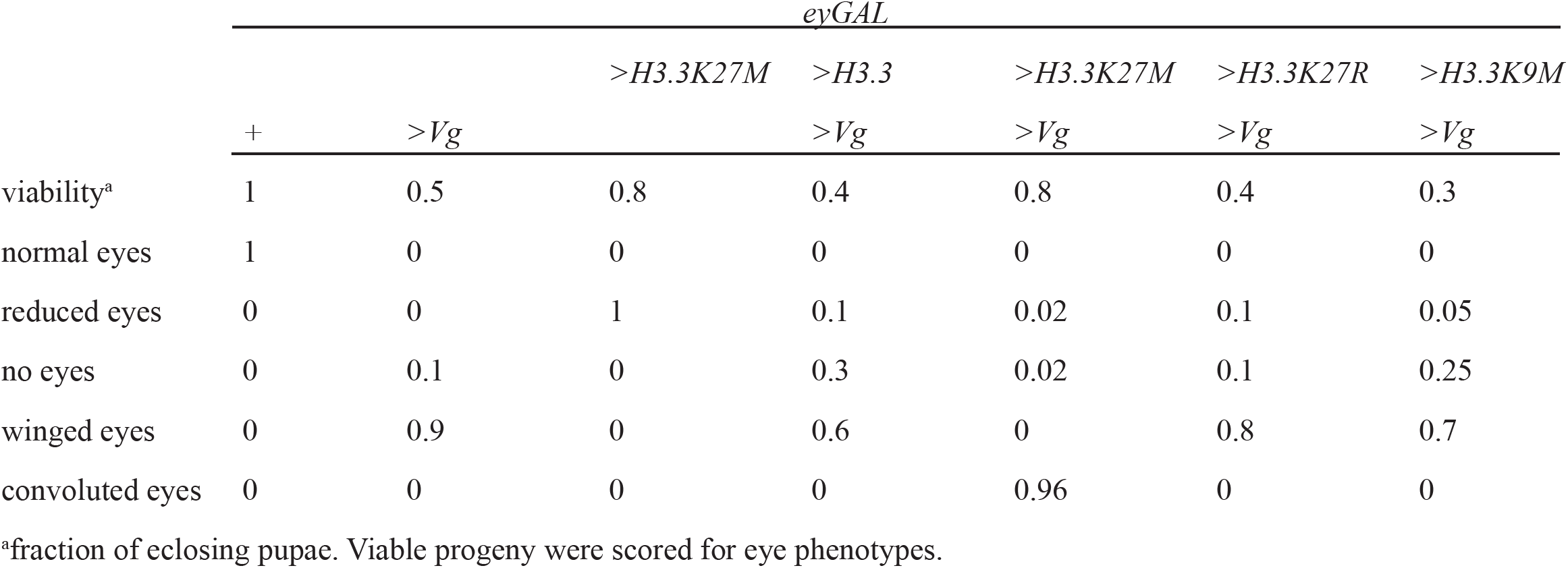
Effect of induced Vg and H3.3 mutations on eye-to-wing reprogramming.

**Figure 1.**
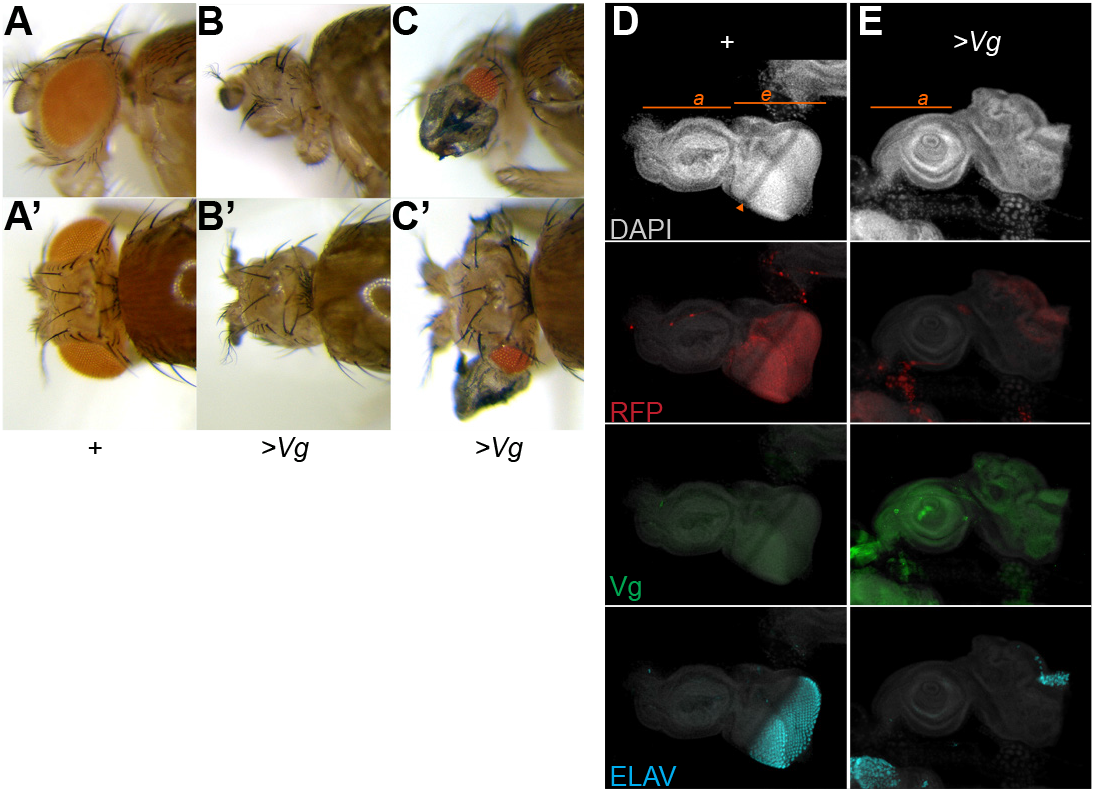
Vg expression transforms the eye into wing tissue. (A-C) Side and dorsal views of the head and eyes of wildtype adults (A) and *eyGAL* >*Vg* adults (B,C). Most adults with >*Vg* expression display pigmented wing-like projections, while some lack eyes entirely. (D,E) Eye-antennal imaginal discs from wildtype larvae (D) and from larvae with >*Vg* expression (E). The *eyGAL* driver also induces a *UAS-RFP* reporter, and discs were immunostained for the Vg protein and for ELAV, a marker of differentiating photoreceptors. The antennal (a) and eye (e) portions of the disc are indicated, and the morphogenetic furrow that separates mitotically active undifferentiated cells in the anterior of the eye from differentiating photoreceptors in the posterior is marked with an orange arrowhead. The eye portions of discs with Vg expression are distorted with low RFP expression, Vg expression, and very few differentiating photoreceptors at the posterior edge.

We dissected and immunostained developing eye-antennal imaginal discs to determine the cellular effects of ectopic Vg and H3.3K27M. The *eyGAL* driver induces expression of an *RFP* reporter gene, marking the eye anlagen of the imaginal disc (Figure 1D). Inducing Vg expression reduces and distorts the eye portion of discs, with very few photoreceptors developing at the posterior edge of the eye disc (Figure 1E). RFP reporter expression is also reduced in these discs, consistent with the inactivation of the *eyGAL* driver as Vg reprograms the eye disc.

We used a transgene encoding an inducible *H3.3K27M* gene [Ahmad & Spens 2019] and the same *eyGAL* driver to express the oncohistone in the eye. Oncohistone expression decreases the size of the adult eye in proportion to the amount of oncohistone produced (Figure 2A-C). Thus, while the oncohistone is associated with proliferation in cancers, expression on its own inhibits tissue growth. Expression of the oncohistone results in reduced staining for H3K27me3 specifically in the eye portion of the disc with normal levels of staining in the antennal portion (Figure 2D,F,G). These discs are slightly smaller than wildtype discs, but with normal morphology and development of photoreceptors (Figure 2D,G). In contrast, the H3K27me3 modification is unaffected by ectopic expression of Vg, with high levels in cells of both the eye and antennal portions of discs (Figure 2H).

**Figure 2.**
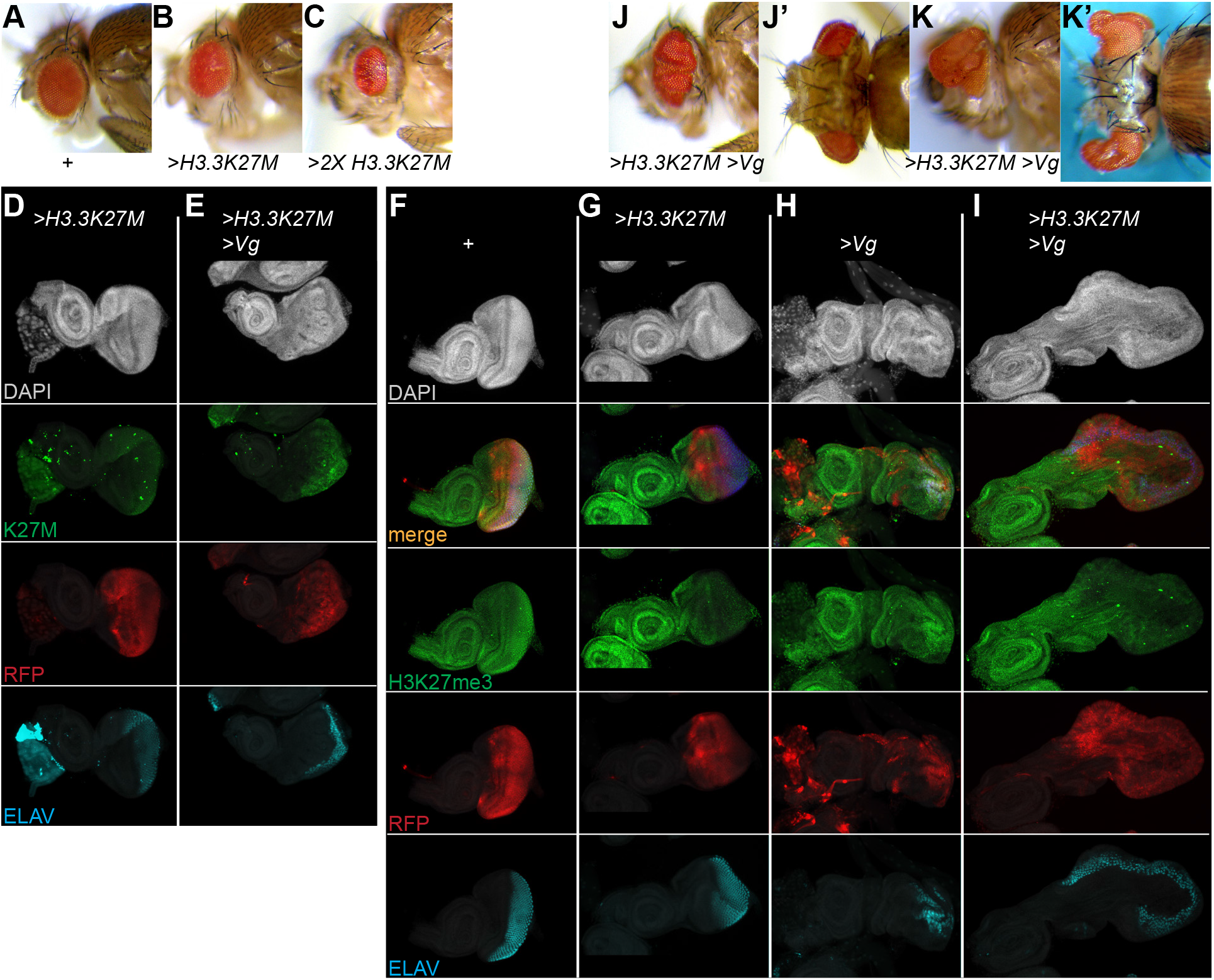
H3.3K27M oncohistones block direct reprogramming. (**A-C**) Side views of eyes of wildtype adults (**A**), adults with one *H3.3K27M* transgene (**B**), or with two *H3.3K27M* transgenes (**C**). Expression of the oncohistone reduces the size of the eye. (**D,E**) Eye-antennal imaginal discs from larvae with *eyGAL*-induced expression of *RFP*, *H3.3K27M*, and *Vg*. Discs were immunostained for the K27M epitope and for the photoreceptor ELAV marker. The oncohistone is expressed throughout the developing eye portion of the disc. (**F-I**) Eye-antennal imaginal discs with *eyGAL*-induced expression of *RFP*, *H3.3K27M*, and *Vg*, and immunostained for the H3K27me3 histone modification and for ELAV. Histone methylation is reduced in the eye portion of the disc relative to staining in the antennal portion in discs expressing the oncohistone. (*J,K*) Examples of adults with co-expression of Vg and H3.3K27M in the eye. Eyes are overgrown and convoluted (J) or overgrown with extreme projections (K).

To test the importance of H3K27me3 chromatin silencing in reprogramming, we co-induced expression of Vg and the K27M oncohistone together in developing eyes. Surprisingly, co-expression completely suppressed eye-to-wing reprogramming (Table 1). The majority of animals co-expressing Vg and H3.3K27M eclose, but have grossly disrupted eyes with a variety of projections, stalks, and folds (Figure 2J,K). These convoluted eyes contain ommatidia with no wing-like tissue, suggesting that co-expression of the oncohistone completely blocks the reprogramming effects of Vg, thereby maintaining the eye identity of these cells. This is apparent in developing imaginal discs, where the eye portion of discs is greatly expanded where the oncohistone is expressed, with developing photoreceptors along a convoluted edge of the disc (Figure 2E,I). Suppression of reprogramming is specific to the K27M oncohistone, because a transgene with a wildtype H3.3 histone, with an H3.3K27R substitution, or with an H3K9M substitution [Herz et al, 2014] do not block reprogramming (Table 1). We tested additional independent *eyGAL4* driver constructs and insertions to induce the H3.3K27M oncohistone and to induce Vg (see Key Resources Table), all of which reproduced the phenotypes of Vg-induced reprogramming and oncohistone-induced overgrowth. Thus, transcription factor-induced reprogramming of the eye specifically requires H3K27me3-mediated silencing.

### Vg and H3K27M induce cell death and proliferation

The sizes of imaginal discs and adult eyes are consistent with decreased proliferation with either Vg or with H3.3K27M expression, but increased proliferation with the two proteins co-expressed. We examined cell division rates and cell death rates by staining eye imaginal discs with a mitotic marker histone (H3S10-phosphorylation) and with a cell death marker (cleaved DCP-1). In wildtype controls, mitoses are scattered throughout the anterior portion of the eye disc, but mostly absent in the posterior region once photoreceptors start to differentiate (Figure 3A) with negligible cell death (Figure 3E). Expression of H3.3K27M does not affect mitosis in the disc (Figure 3B), but a stripe of cell death appears across the disc where cells transition from proliferating to differentiating regions (Figure 3F). This implies that the reduction in eye size with H3.3K27M expression is at least in part due to reduced cell viability. In contrast, induction of Vg is associated with increased mitosis and increased cell death throughout the disc (Figure 3C,G). Thus, reduced cell viability limits the size of the reprogrammed tissue. Finally, co-expression of H3.3K27M and Vg produces very large discs with patches of mitosis frequently apparent (Figure 3D), but these discs show extensive cell death in undifferentiated regions (Figure 3H). The hyperproliferation of some regions accounts for overgrowth in spite of extensive oncohistone-induced cell death in other regions.

**Figure 3.**
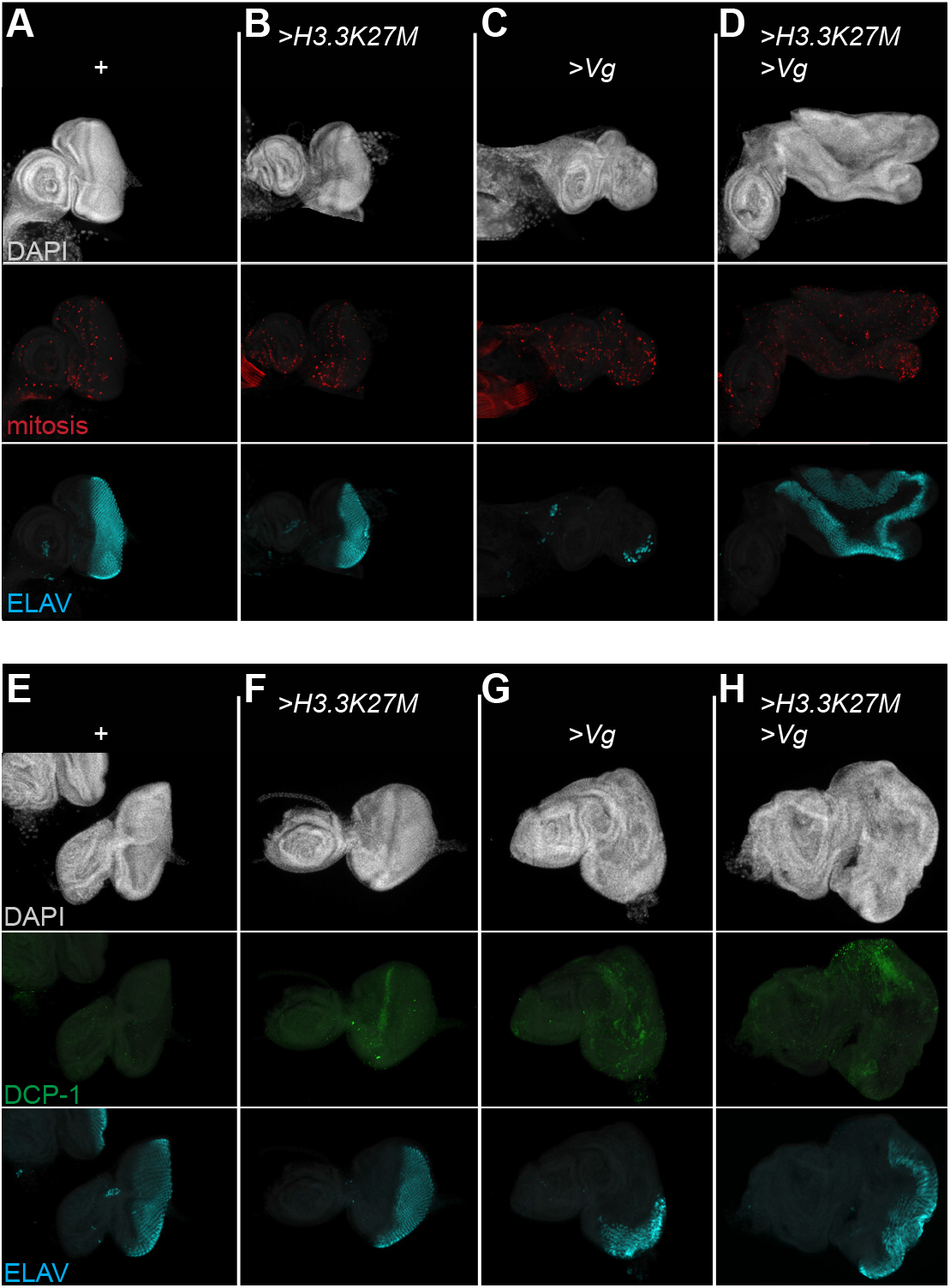
H3.3K27M and Vg stimulate cell death and proliferation. Eye-antennal discs with *eyGAL*-induced expression of H3.3K27M and Vg and immunostained for the H3S10-phosphorylation marker of mitosis (**A-D**) and for the cleaved DCP-1 marker of apoptosis (**E-H**). All discs were immunostained for ELAV.

### Gene expression in Vg-reprogrammed tissues

Reprogramming of the eye inactivates the *eyGAL* driver once cells transform, and this is apparent by reduced RFP expression when Vg is induced (Figure 1E). Inactivation of the *eyGAL* driver implies that the Vg transgene will also be inactivated, and so the endogenous *vg* gene must be activated for successful reprogramming. To test this, we constructed animals with the *eyGAL* driver and the inducible Vg transgene, but lacking the endogenous *vg* gene. As expected, these animals have very small eyes with no wing tissue outgrowths (Figure 4A). This implies that transient expression of Vg activates endogenous genes for wing specification. The endogenous *vg* gene is also required for overgrowth of eye discs with oncohistone and ectopic Vg expression, as a dominant negative *vg^U^* allele reduces eye size (Figure 4B).

**Figure 4.**
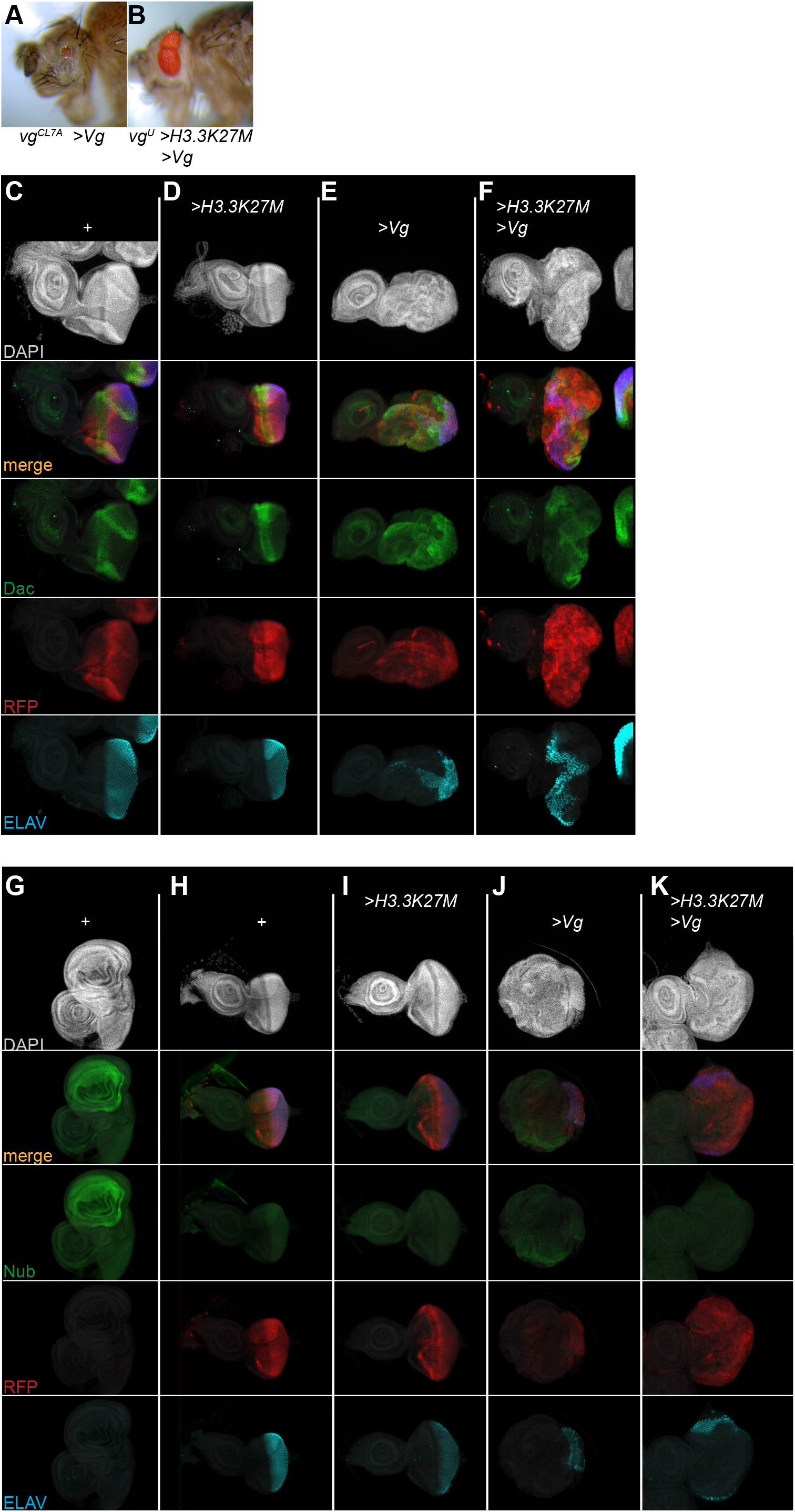
Tissue-specific gene expression changes during direct reprogramming. **A**) Side view of a *vg*^*CL7A*^ mutant adult with *eyGAL*-induced Vg expression in the eye. Eye size is reduced, but not transformed. (**B**) Side view of a *vg*^*U*/+^ head with *eyGAL*-induced H3.3K27M and Vg co-expression in the eye. The eye is convoluted, but reduced in size instead of overgrown. (**C-F**) Eye-antennal discs with *eyGAL*-induced expression of RFP, H3.3K27M and Vg and immunostained for the eye-specific Dac and ELAV proteins. Neither Vg nor H3.3K27M expression prevent Dac expression, although these discs are disorganized. (**G-K**) Wildtype wing imaginal discs (**G**) and eye-antennal discs with *eyGAL*-induced expression of H3.3K27M and Vg (**H-K**) immunostained for the wing-specific Nub protein and ELAV. The wing pouch is heavily labeled with Nub, and eye discs with Vg expression have patches with low level Nub staining.

To characterize activation of wing-specific genes further, we stained discs for proteins specific for eyes or for wings. Expression of the wing specification factors Nubbin (Nub) in reprogrammed eye discs is weaker than in wing discs (Figure 4G-I), implying that the wing determination program is not efficiently activated. These discs continue to express the eye factor Dacshund (Dac), although the normal striped pattern of this factor is distorted by the gross disorganization of the reprogrammed disc (Figure 4C-F). Notably, there is no expression of Nub in >*H3.3K27M* >*Vg* overgrown discs (Figure 4J), while RFP and Dac are highly expressed. Thus, in this setting the ectopic expression of Vg does not activate wing determination and eye factors continue to be expressed (Figure 4F).

### Chromatin profiling of H3K27me3 domains in reprogrammed tissues

To profile chromatin domains in Vg-reprogrammed eyes, we adapted the CUT&Tag method [Kaya-Okur et al, 2019] for dissected imaginal discs from Drosophila larvae. CUT&Tag works by first soaking unfixed cells with a factor-specific antibody which binds to chromatin sites, followed by decoration with a secondary antibody. Next, a protein-A-Tn5 (pA-Tn5) transpososome loaded with sequencing adapters is soaked in, binding to the chromatin-bound antibodies. Activation of the tethered transpososome by adding magnesium then integrates sequencing adapters around binding sites, and sequencing of the resulting library thus maps the targeted chromatin protein. We have previously adapted the micrococcal nuclease-based CUT&RUN method for dissected imaginal discs [Ahmad & Spens, 2019], and adjusting buffers for CUT&Tag works reliably. For CUT&Tag with tissue samples, we dissect wing or eye imaginal discs, coat them with Concanavalin A magnetic beads for handling, lightly permeabilize them with digitonin, and sequentially incubate with antibodies and then with pA-Tn5. Resulting libraries are subjected to Illumina paired-end sequencing and mapped to the Drosophila dm6 genome assembly. Imaginal discs from 2-3 larvae were sufficient to generate chromatin profiles, although the capacity of CUT&Tag to profile very small sample sizes should work with even less tissue [Kaya-Okur et al, 2019; Kaya-Okur et al, 2020].

We first mapped the H3K27me3 silencing modification in eye and wing imaginal discs from wildtype larvae (Supplementary Information; Supplementary Figure 1A-C). Previous studies found that H3K27me3-marked domains are shared between larval tissues but with quantitative differences in chromatin methylation that correspond to expression of included genes [Loubière et al, 2016; Ahmad & Spens, 2019]. For example, a 350 kb H3K27me3 domain encompasses the *ANTENNAPEDIA-COMPLEX* (*ANTP-C*) cluster of homeobox genes in both eye and wing imaginal disc cells (Figure 5A). In eye imaginal discs the *Antp* gene is silenced and heavily coated with H3K27me3-marked chromatin, while in wing imaginal discs the *Antp* gene is transcribed, and H3K27me3 across this gene is correspondingly depleted. Similarly, H3K27me3 domains with regulatory genes differentially-expressed between wings and eyes show quantitative changes in histone methylation: chromatin encompassing the wing-specific *apterous* (*ap*) gene is methylated in both eye and wing imaginal discs, but noticeably reduced in wing tissue, while the eye-specific *sine oculus* (*so*) and *optix* genes have less methylation in eye tissue (Figure 5B). Thus, activation of tissue-specific genes in H3K27me3 domains is accompanied by the reduction of histone methylation, but complete removal of this mark is not required for expression.

**Figure 5.**
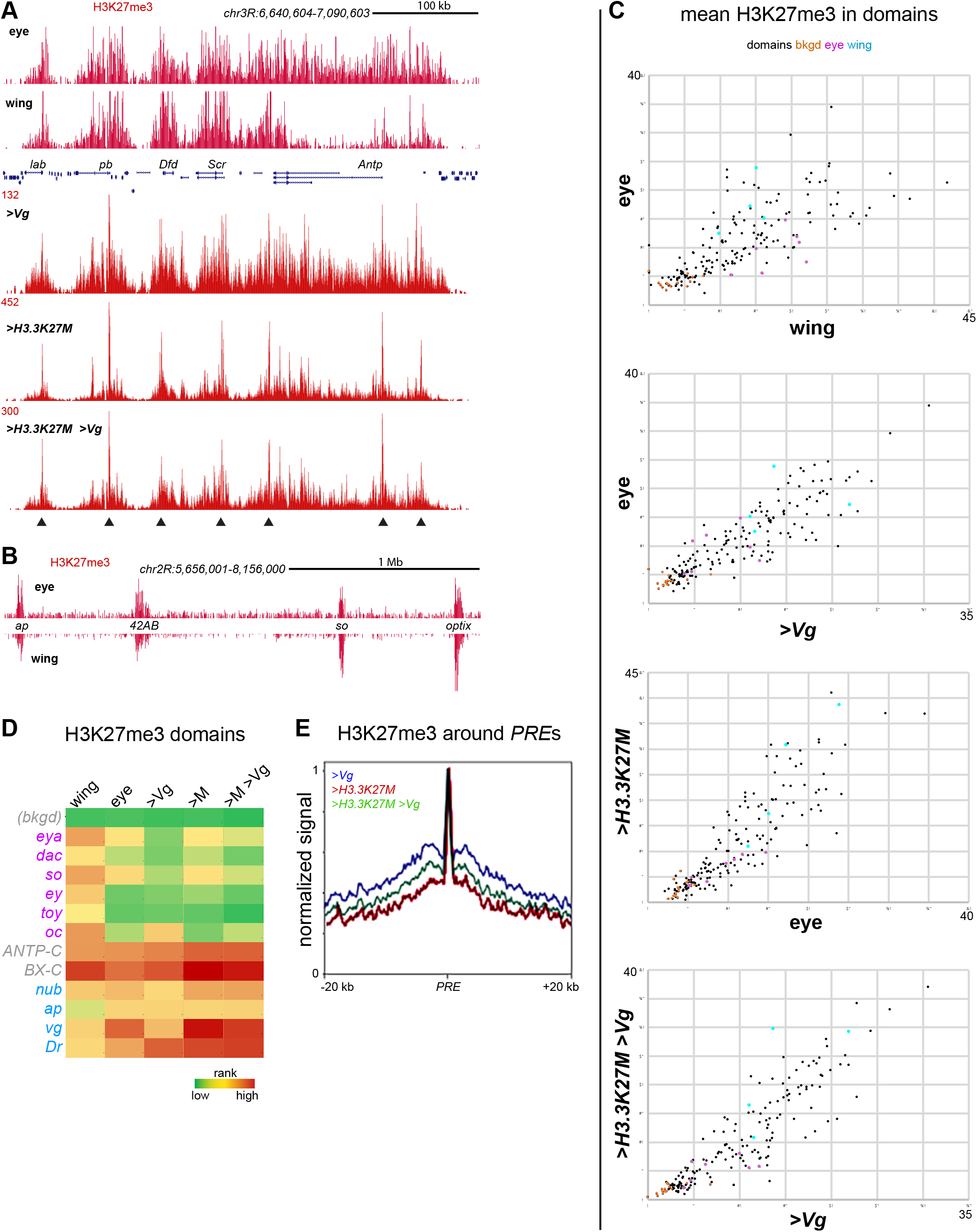
Chromatin profiling of H3K27me3 domains in reprogrammed eye imaginal discs. (**A**) Chromatin profiling at the *ANTP-C* H3K27me3 domain in wing and eye imaginal discs. Arrowheads mark the positions of major Polycomb-bound *PRE*s. (**B**) H3K27me3 profiling spanning the wing-specific *ap* and eye-specific *so* and *optix* genes in wing and eye imaginal discs. (C) Scatterplots of average H3K27me3 signal in 166 domains (black) and 16 background control regions (orange). Domains including eye specification genes are in magenta, and domains with wing specification genes are in blue. (D) Domains ranked by decreasing average H3K27me3 signal in wing and eye imaginal discs. Selected domains spanning eye (magenta), wing (blue) or common domains (grey) are displayed. (E) Average H3K27me3 signal around 700 Polycomb-bound *PRE*s defined from published ChIP-seq profiling [Loubière et al, 2016] within H3K27me3 domains.

To quantify changes in chromatin landscapes, we measured the average H3K27me3 signal across 166 annotated H3K27me3 domains and 16 control regions (Supplementary Information) in wing and eye discs and compared these on a scatter plot (Figure 5C). Domains that encompass genes involved in eye specification have lower chromatin methylation in eye discs and gain signal in wing discs when the genes are inactive. The opposite trend occurs for four wing specification genes. Comparing the rank order of average domain methylation, eye specification genes substantially lose methylation (Figure 5D). In contrast, wing specification genes gain methylation in eye tissue, even though 3 of these 4 genes have substantial H3K27me3 signal in both tissues (Figure 5D).

We dissected away the antennal portion of the disc to profile H3K27me3 in reprogrammed eye tissue without contamination from wildtype antennal cells. Unlike the differences between eye and wing discs, expression of Vg in eye discs results in little change in H3K27me3 across domains compared to their chromatin signal in eye discs (Figure 5C,D; Supplementary Figure 1D). Thus, reprogramming of eye cells is not accompanied by dramatic changes in chromatin methylation patterns.

We next looked at the effect of the H3.3K27M oncohistone on H3K27me3 patterns in the eye. The oncohistone is incorporated into chromatin by histone H3.3-specific chaperones at active promoters and enhancers, and at lower levels by DNA replication throughout the genome [Chan et al, 2013; Piunti et al, 2017; Mohammad et al, 2017; Sarthy et al, 2020], and expression of H3.3K27M in eye discs gives a similar promoter enrichment with background throughout the genome when profiled using an antibody to the K27M epitope (Supplementary Figure 1E). Expression of the oncohistone reduces H3K27me3 staining (Figure 2G,I), and quantitative chromatin profiling of glioma cell lines has previously shown that H3K27me3 is globally reduced but a few domains remain [Sarthy et al, 2020]. By profiling eye portions of imaginal discs with oncohistone expression, we find that H3K27me3 still coats domains, although with more dispersion between domains that we attribute to the lower H3K27me3 signal in oncohistone-expressing cells (Figure 5C). In Drosophila H3K27me3 domains are nucleated at nucleosome-depleted *Polycomb Response Elements* (*PRE*s), and chromatin methylation then spreads out across the domain [Laprell et al 2017]. The H3.3K27M oncohistone specifically inter feres with methylation spreading [Harutyunyan et al, 2019; Jain et al, 2020]. Indeed, changes in the pattern of H3K27me3 methylation is apparent at individual domains, such as the *ANTP-C* (Figure 5A). Here, *PRE*s are still heavily marked with the histone modification, but intervening regions are reduced, consistent with the oncohistone inhibiting spreading of the modification from *PRE*s. We used published sites of Polycomb binding within H3K27me3 domains [Loubière et al, 2016; Supplementary Information] to define the positions of *PRE*s and display H3K27me3 signals around those sites, confirming a consistent loss of methylation surrounding a peak immediately over *PRE*s (Figure 5E). H3K27me3 profiling of eye disc portions co-expressing Vg and H3.3K27M recapitulate this effect: domains are still present but show reduced spreading of H3K27me3 around *PRE*s (Figure 5E). The one notable change in H3K27 methylation is at the *vg* gene: ectopic Vg expression reduces methylation across this domain, but this gene becomes more methylated with oncohistone co-expression (Figure 5D). Thus, oncohistone expression in the eye does not ablate H3K27me3 domains, instead, the genome-wide reduction in methylation affects both eye- and wing-specific domains.

### Chromatin profiling of H3K4me2 histone modification in reprogrammed tissues

We next looked at chromatin profiles of the H3K4me2 histone modification, a marker of active promoters and enhancers. The genome-wide enrichment of H3K4me2 at promoters is very similar between wing and eye discs, driven by their similar expression profiles of housekeeping genes (Figure 6A). However, promoters of wing and eye specification transcription factors do show increased H3K4me2 signal in the tissue where the gene is active (Figure 6B,C). H3K4me2 signal also identifies which promoter is active between alternative promoters, such as at the *ey* gene (Figure 6C). We used published mRNA profiling of wing and eye imaginal discs [Naval-Sanchez et al, 2013] to de fine differentially-transcribed genes between these tissues, identifying 804 genes up-regulated in eye discs and 174 up-regulated in wing discs, and rank-ordered the amount of H3K4me2 signal in a 1 kb window centered on annotated gene promoters in the Drosophila genome (Supplementary Information). In most cases promoters gain H3K4me2 signal in the tissue where they are active, and the few exceptions appear to be the unused promoter of alternative TSSs (Figure 6D). Thus, the activation of specification genes is associated with both increases in promoter H3K4me2 signal reductions in domain H3K27me3 signals.

**Figure 6.**
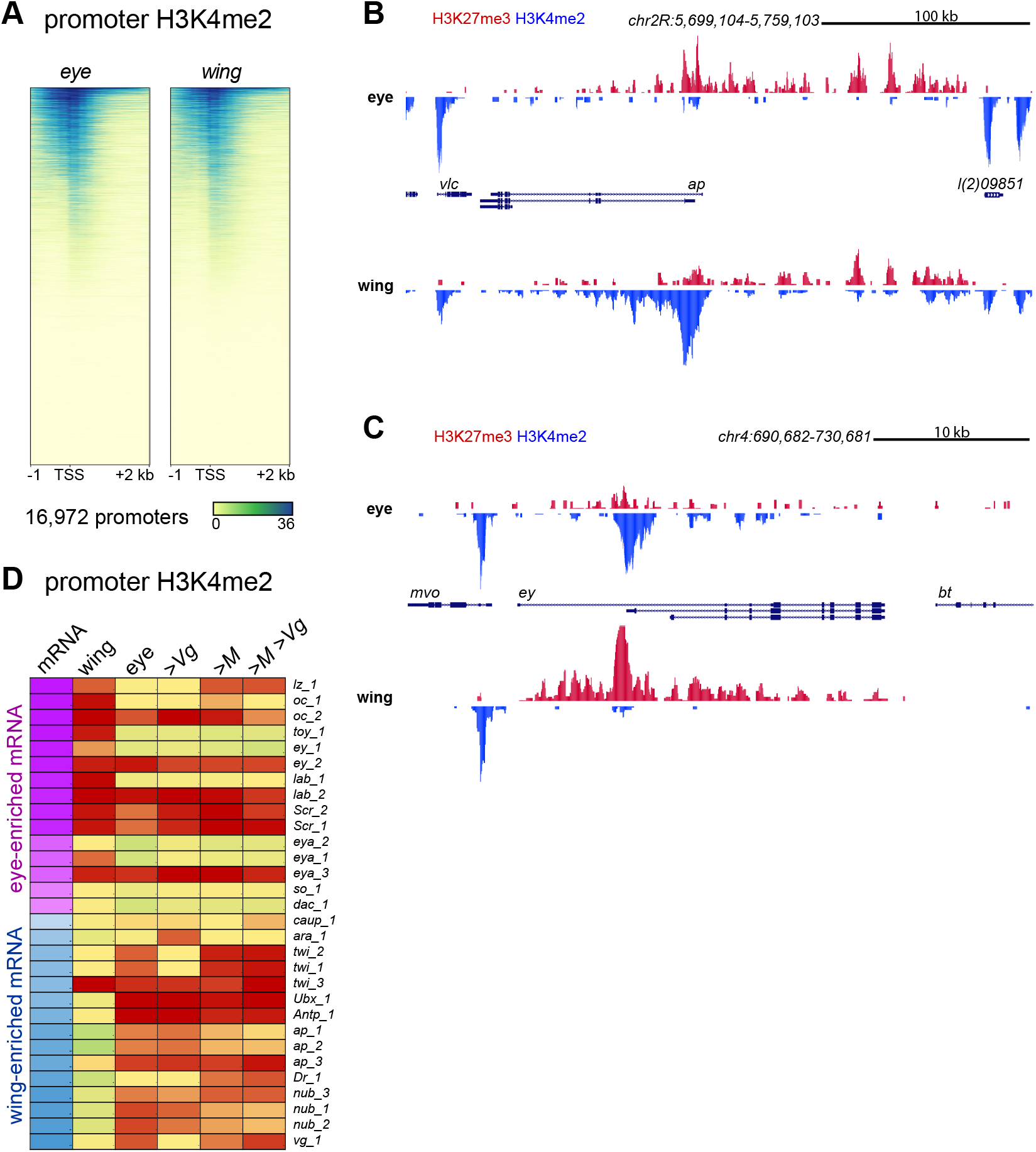
Chromatin profiling of H3K4me2 signal at promoters in reprogrammed eye imaginal discs. (**A**) Heatmaps of H3K4me2 signal around promoters in eye and wing imaginal discs. (**B,C**) H3K27me3 and H3K4me2 features at the wing-specific *ap* gene (**B**) and at the eye-specific *ey* gene (**C**) in wing and eye imaginal discs. (**D**) Promoters ranked by decreasing average H3K4me2 signal in a +500 bp around the TSS in wing and eye imaginal discs. Selected promoters of differentially-expressed genes defined from published RNA-seq profiling [Naval-Sanchez et al, 2013] are displayed. Alternative promoters for these genes are numbered.

We then profiled the H3K4me2 modification in dissected eye discs from larvae expressing Vg. Promoter signals in these discs show slight reduction in H3K4me2 at the promoters of multiple eye specification genes such as *oc*, *Scr*, *ey*, *eya*, and *so* (Figure 6D). These changes are not as dramatic as the loss of H3K4me2 signal in wing discs, perhaps due to the mosaicism of this sample (Figure 1E) or to incomplete activation in transformed cells. However, some wing-specific genes in Vg-expressing eye discs do show gain in H3K4me2 signal at gene promoters, such as at twi and at vg itself (Figure 6D). These limited promoter changes indicate that reprogramming of the eye by ectopic Vg is driven by the activation of a subset of genes involved in wing specification.

We then profiled H3K4me2 histone modifications in eye discs expressing the H3.3K27M oncohistone. Expression of the oncohistone has little effect on H3K4me2 marks at active promoters in eye discs (Figure 6D; Supplementary Figure 1F). Co-expression of H3.3K27M and Vg also has little effect on promoter H3K4me2 marks, which is expected since ectopic Vg expression also has limited effects. The largest change is at the *vg* promoter itself, where ectopic Vg expression increases H3K4me2 signal at this promoter, but co-expression of Vg and H3.3K27M prevent this gain (Figure 6D). Ectopic Vg expression also causes slight reductions of H3K4me2 signal at the *ey* and *eya* promoters, and these signals increase when Vg and the oncohistone are co-expressed (Figure 6D). These changes are consistent with the idea that Vg expression inhibits expression of eye specification genes during reprogramming, but oncohistone co-expression relieves this inhibition.

### H3K27M and Vg co-expression imitates PRC1 mutants

Tissue reprogramming has been associated with a number of components in the Polycomb silencing system [Grossniklaus & Paro, 2014]. H3K27 trimethylation is catalyzed by the E(z) histone modifying enzyme, a component of the Polycomb Repressive Complex 2, while the Polycomb Repressive Complex 1 (PRC1) complex binds the H3K27me3 mark on nucleosomes [Di Croce & Helin, 2013]. Mutation of PRC1 components result in overgrowth of imaginal discs, but mutations in PRC2 components inhibit cell growth [Classen et al, 2009; Loubière et al, 2016], implying that the two complexes have non-overlapping functions. We examined the roles of PRC1 and PRC2 in reprogramming of the eye. A previous study characterized *eyGAL*-induced knock-down of PRC1 and PRC2 components [Zhu et al, 2018], which we repeated here. Knocking down the PRC2 histone methyltransferase E(z) inhibits proliferation, resulting in near elimination of the eye portion of imaginal discs (Figure 7A) and lethality, where dying animals lack all head structures derived from the eye-antennal imaginal disc (Figure 7C). Thus, growth inhibition by H3.3K27M expression resembles intermediate reduction of the E(z) enzyme, consistent with the dominant inhibition of this enzyme by the oncohistone.

**Figure 7.**
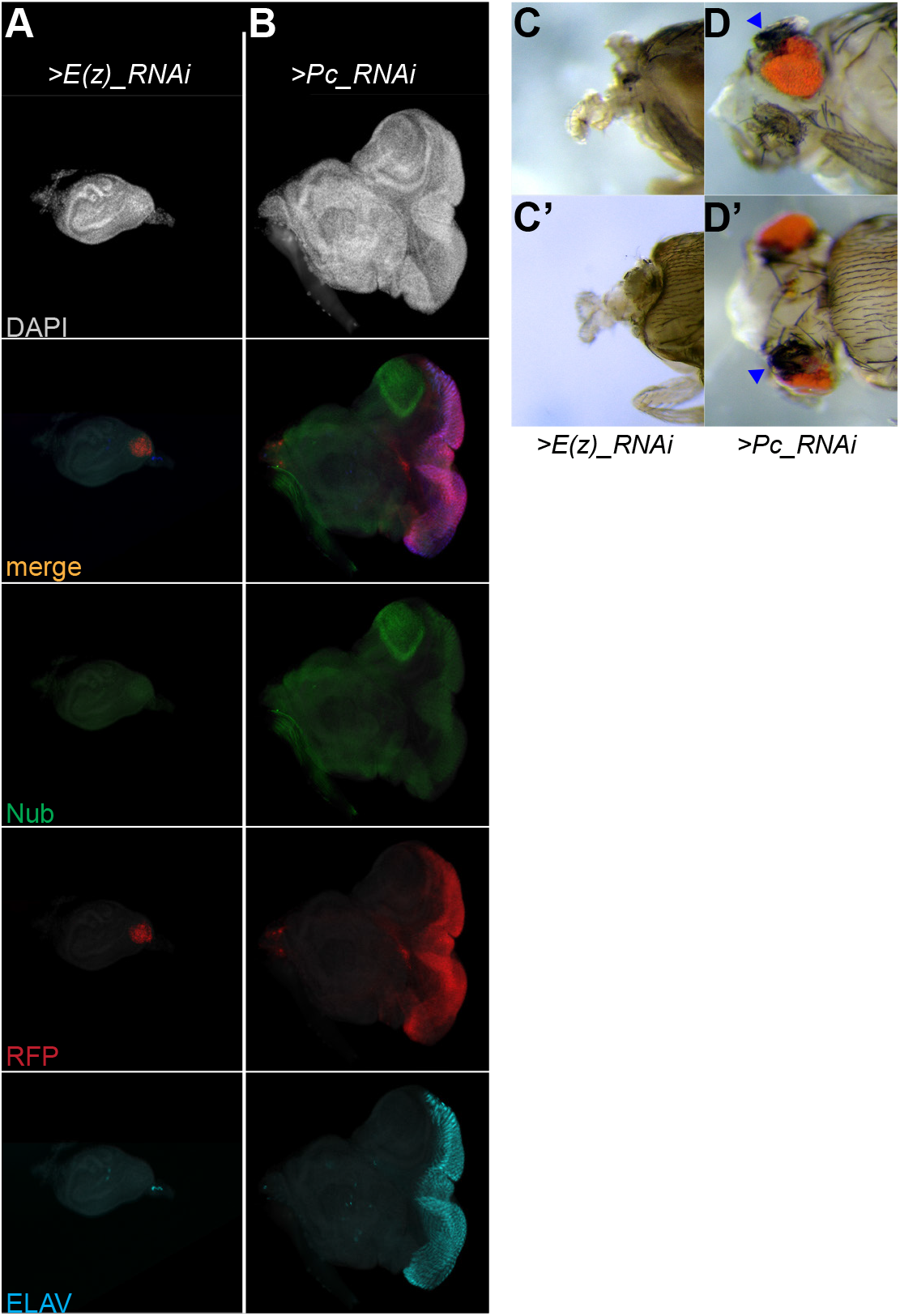
Effects of reduced PRC1 and PRC2 on direct reprogramming and cell proliferation. (**A,B**) Eye-antennal discs with *eyGAL*-induced expression of RFP and RNAi constructs directed against *E(z)* (**A**) or *Pc* (**B**) transcripts. Both discs are immunostained for the wing-specific Nub protein and the photoreceptor marker ELAV. The eye and antennal portions of discs are greatly reduced with *E(z)* knockdown, while *Pc* knockdown results in overgrowth of the eye portion and expression of Nub at the dorsal edge of the disc. (**C,D**) Heads of dissected dying pharate pupae with *E(z)* knockdown (**C**) or *Pc* knockdown (**D**). Loss of *E(z)* ablates all tissue derived from the eye-antennal disc including the head capsule and eyes, without affecting mouthparts. Loss of *Pc* induces ectopic wing-like structures (blue arrowheads) at the dorsal edges of eyes.

Eye-specific knockdown of Polycomb derepresses the *Antp* HOX gene, reprogramming part of the eye to wing-like tissue and inducing overgrowth in the disc [Zhu et al, 2018]. This is also lethal, however knockdown eye discs are often expanded with an outgrowth on one side of the disc [Zhu et al, 2018] and (Figure 7B). Dying animals with Polycomb knockdown show wing-like outgrowths on the dorsal side of the eye [Zhu et al, 2018] (Figure 7D). Polycomb loss efficiently induces wing tissue, since the wing factor Nub is expressed at high levels in these discs (Figure 7B). The location of high Nub expression coincides with a region that appears to be predisposed to transformation by high expression of common signalling factors [Maves & Schubiger, 1999; Zhu et al, 2018]. Thus, effects of PRC1 loss actually resemble those of Vg and H3.3K27M co-expression, implying that the loss of chromatin silencing and mis-expression of a determinative transcription factors synergizes to drive tissue overgrowth.

## Discussion

The idea that cell fate programs are reinforced by chromatin silencing of alternative pathways implies that reducing epigenetic barriers that restrict cell fates will stimulate cell fate transformations. We find the opposite – using ectopic expression of the Vg master regulator factor, we find that compromised silencing does not enhance transformation of the eye; instead cells of the eye disc hyperproliferate. It is startlingly simple to create overgrowth tumors in the Drosophila developing eye: expression of only two proteins – the H3.3K27M oncohistone and a transcription factor – are sufficient. Commitment to a developmental fate requires both the expression of genes for determinative transcription factors and the silencing of genes for alternative fates. Coordinated activation and silencing may shape and stabilize developmental trajectories. Our results highlight that chromatin silencing is essential for transcription factor-induced developmental reprogramming. Cell identity in the Drosophila eye is determined by a network of self-reinforcing transcription factors. To reprogram this tissue expression of Vg must both induce wing specification genes and silence eye-specifying factors. Surprisingly, inhibition of H3K27 methylation does not enhance reprogramming, implying that the activation of silenced wing specification genes is not limiting. Instead, our results argue that H3K27me3-mediated silencing of eye-specific factors is needed for successful reprogramming. This requirement may result from the inducing Vg in a setting where the eye determination program has already been established. Further, the differentiation of retinal cells even when Vg is expressed indicates that eye determination factors must be dominant to wing specification factors, similar to what has been observed after tissue damage in Drosophila [Maves & Schubiger, 2003]. Intriguingly, overexpression of the histone gene transcription factor Wge can also drive eye-to-wing reprogramming, but this case requires chromatin silencing mediated by the histone H3K9 methyltransferase Su(var)3-9 [Ozawa et al, 2016; Masuko et al, 2018]. Thus, an established developmental program may be silenced by either H3K27me3- or H3K9me3-mediated pathways during reprogramming.

Why does inhibiting reprogramming with the H3K27M oncohistone result in neoplastic growth? Genetic studies in mammalian systems have demonstrated that H3K27M oncohistones are not sufficient to induce tumorigenesis on their own [Mohammad et al, 2017]. Instead, secondary mutations are necessary, and the low mutational burdens of H3K27M-bearing cancers in patients implies that specific mutations are sufficient for malignancy. Some mutations are in classical tumor suppressor genes like *TP53*, thereby enhancing malignancy. In pediatric midline gliomas, additional secondary mutations are often in specific signaling receptors such as *PDGFRA* or *ACVR1* [Schwartzentruber et al, 2012; Fontebasso et al, 2014; Buczkowicz et al, 2014; Mackay et al, 2017; Wu et al, 2014], and the hyperactivation of developmental signalling probably induces the mis-expression of developmental transcription factors. Co-expression of the H3.3K27M oncohistone and Vg appears to imitate this effect. Vg both promotes a wing cell fate and stimulates cell growth [Delanoue et al, 2004], thus suppressing reprogramming may result in unrestrained proliferation. While mutation of the four *Vg* homologs in humans is not observed in gliomas, mis-expression of three paralogs are associated with other cancers [Yamaguchi, 2020]. More generally, that simply mis-expressing a developmental transcription factor and an oncohistone stimulates proliferation suggests that other transcription factors might also do so, explaining the cell-type- and stage-restriction of oncohistone-driven cancers.

It is striking that mutations in PRC1 components stimulate uncontrolled proliferation, while mutation of PRC2 components do not [Classen et al, 2009; Loubière et al, 2016]. These two complexes normally work together to silence domains in genomes, but PRC1 also regulates gene expression of some developmental targets [Aranda et al, 2015; Loubière et al, 2020]. Our results suggest that mutations in PRC1 components both lose chromatin silencing and mis-express a developmental regulator, driving cell proliferation.

## Methods

### Fly strains

All crosses were performed at 25°C. All mutations and chromosomal rearrangements used here are described in Flybase (http://www.flybase.org) and sources are listed in the Key Resources Table. The *eyGAL4*-3-8 driver was used for all experiments shown here, although identical results were obtained with other *eyGAL4* constructs and insertions.

### Transgenes

Inducible H3.3 constructs were constructed by cloning the *UASp* promoter and the *His3.3A* ORF into the pKC27mw transformation vector [Okluski et al, 2011] (a gift from L. Ringrose, Humboldt-Univerität zu Berlin), and then used site-directed mutagenesis PCR to make the K27M and K27R mutated versions. Each plasmid was injected into *y M[vas-int.Dm]*ZH-2A *w; P[attP,y+,w3’]*VIE-260B embryos by Bestgene Inc (Chino Hills, CA). This line contains two landing sites [Green et al, 2014]; integrants at the 25C landing site were used in this study.

### Imaging imaginal discs

Imaginal discs from late 3rd instar larvae were dissected and fixed for 10 minutes in 4% formaldehyde/PBST (PBS with 0.1% triton-X100), and then incubated twice in 0.3% sodium deoxycholate/PBST for 20’ each. Samples were incubated with primary antiserum diluted in PBST supplemented with 10% goat serum at 4° overnight, and finally with fluorescently labeled secondary antibodies (1:200 dilution, Jackson ImmunoResearch). All tissues were stained with 0.5 μg/mL DAPI/PBS, mounted in 80% glycerol on slides, and imaged by epifluorescence on an EVOS FL Auto 2 inverted microscope (Thermo Fisher Scientific) with a 10X objective. Pseudo-colored images were adjusted and composited in Adobe Photoshop and Adobe Illustrator. All antibodies used are listed in the Key Resources Table.

### Imaging adult eyes

Adults were euthanized in a freezer and then imaged using a Sony digital camera mounted on a Nikon SMZ1500 stereomicroscope. Images were color-corrected with Adobe Photoshop and composited in Adobe Illustrator.

### Chromatin profiling and sequencing

We dissected imaginal discs from 3rd instar larvae in Wash+ buffer (20 mM HEPES pH 7.5, 150 mM NaCl, 0.5 mM spermidine with Roche cOmplete protease inhibitor). We used 4 wing imaginal discs and 6 eye-antennal imaginal discs for each chromatin profiling experiment. Experiments were performed in duplicate and in parallel to minimize technical variation. We used immuno-tethered CUT&Tag chromatin profiling [Kaya-Okur et al, 2020] with antibodies to histone H3K27me3 (C36B11, Cell Signalling Technology) and to histone H3K4me2 (13-0027, Epicypher) modifications. To adapt CUT&Tag for whole tissues, we coated imaginal discs with BioMag Plus Concanavalin-A-conjugated magnetic beads (ConA beads, Polysciences, Inc) in 8-tube PCR strips, and exchanged solutions on a magnetic stand (MSR812, Permagen). Tissues were incubated with primary antibody in dbe+ buffer (20 mM HEPES pH 7.5, 150 mM NaCl, 0.5 mM spermidine, 2 mM EDTA, 1% BSA, 0.05% digitonin with Roche cOmplete protease inhibitor) overnight at 4°C, incubated with secondary antibody in dbe+ buffer for 1 hour at room temperature, and then incubated with protein-A-Tn5 loaded with adapters in 300Wash+ buffer (20 mM HEPES pH 7.5, 300 mM NaCl, 0.5 mM spermidine with Roche cOmplete protease inhibitor) for 1 hour. After one wash with 300Wash+ buffer, samples were incubated in 300Wash+ buffer supplemented with 10 mM MgCl2 for 1 hour at 37° to tagment chromatin. Reactions were stopped by addition of SDS to 0.16% and protease K to 0.3 mg/mL, incubated at 58° for 1 hour, and DNA was purified by phenol:chloroform extraction and ethanol precipitation.

Libraries were prepared as described [Kaya-Okur et al, 2020], with 14 cycles of PCR with 10 second combined annealing and extension for enrichment of short DNA fragments. Libraries were sequenced for 25 cycles in paired-end mode on the Illumina HiSeq 2500 platform at the Fred Hutchinson Cancer Research Center Genomics Shared Resource. Paired-end reads were mapped to release r6.30 of the D. melanogaster genome obtained from FlyBase using Bowtie2, and to the E coli genome for spike-in normalization. A step-by-step protocol is posted: https://www.protocols.io/view/cut-tag-with-drosophila-tissues-bnx5mfq6

### Data analysis

Track screenshots were produced using the UCSC Genome browser (http://genome.ucsc.edu) [Kent et al, 2002]. We manually annotated H3K27me3 domains as enriched blocks in either wing or eye imaginal discs, or in profiling of larval brains [Ahmad &Spens, 2019], and are listed in Supplementary Information. We used EPDnew for promoter locations in dm6 genome assembly [Dreos et al, 2015], and FB 2020_03 for gene annotations [Gramates et al, 2017].

Analysis and display was done using deeptools in Galaxy (multibigwigsummary, computematrix, plotheatmap, plotPCA) and MS Excel. Differential gene expression and chromatin feature analysis was done using degust (degust.erc.monash.edu).

## Detailed protocol at protocols.io

https://www.protocols.io/view/cut-tag-with-drosophila-tissues-bnx5mfq6

## GEO link

In process

## Acknowledgements

We thank Jay F. Sarthy and Derek H. Janssens for comments. Stocks obtained from the Bloomington Drosophila Stock Center (NIH P40OD018537) and antibodies obtained from the Developmental Studies Hybridoma Bank (created by the NICHD of the NIH) were used in this study. This work was funded by the NIH (R01HG010492, SH).

## Supplementary Materials

### Supplementary Information

Lists of chromatin profiling sites.

### Supplementary Note

Data processing for Figures 5, 6, and Supplementary Figure 1.

#### New data

All sequencing data is deposited in GEO under the super-entry (*in progress*).

#### Published data

EPDnew, FB 20202_3, Naval-Sanchez et al 2013 (GSE59059, wing imaginal disc and GSE59059,eye imaginal disc), Loubiere et al 2016 (GSE74080, Pc profiling in eye imaginal discs).

#### Software packages used

deepTools v. 3.3.2 in usegalaxy.edu

#### Extracting promoter signals

mean signal coverage +500 bp around EPDnew annotated gene TSSs was extracted using ‘multibigwigSummary’ in deepTools. Values for each dataset are provided in Supplementary Information.

#### Extracting H3K27me3 domain signals

mean coverage across annotated H3K27me3 domains was extracted using ‘multibigwigSummary’ in deepTools. Values for each dataset are provided in Supplementary Information.

#### Defining differentially-expressed genes

RNA-seq count tables for wildtype eye and wing imaginal discs [Naval-Sanchez et al, 2013] were displayed in degust v. 4.1.1 (degust.erc.monash.edu), and differentially-expressed genes were selected with a >1.5 fold change cutoff. This list is provided in Supplementary Information.

#### Heatmapping

In all cases heatmaps display −1 kb to +2 kb around annotated TSSs of genes with 10 bp signal binning, ordered by mean signal and plotted using deepTools. The display range was adjusted to the maximum and minimum signals for each heatmap, and the number of regions is reported in the figures.

#### For Figure 5A

UCSC Genome Browser tracks for H3K27me3 (red) datasets 1534 (w eye discs), 1732 (w wing discs), 1805 (>Vg eye discs), 1800 (>H3.3K27M eye discs), 1806 (>H3.3K27M >Vg eye discs) were plotted with autoscaling.

#### For Figure 5B

UCSC Genome Browser tracks for H3K27me3 (red) datasets 1534 (w eye discs) and 1732 (w wing discs) were plotted with autoscaling.

#### For Figure 5C

Mean H3K27me3 signal across annotated H3K27me3 domains and background regions was plotted in MS Excel.

#### For Figure 5D

Mean H3K27me3 signal across annotated H3K27me3 domains and the average of background regions was rank-ordered for each sample and heatmapped using three-color conditional formatting in MS Excel with Green-Yellow-Red for low-to-high values. Selected values are shown in the figure. The full domain list is provided in Supplementary Information.

#### For Figure 5E

We used 2000 called Polycomb-bound sites in eye imaginal discs [Loubiere et al, 2016] and selected the top 700 sites that fall within H3K27me3-marked domains. This list is provided in Supplementary Information. We extracted and displayed mean H3K27me3 signal +20 kb around these sites using the ‘computematrix’ and ‘plotHeatmap’ functions of deepTools with 10 bp binning. Plots were individually scaled to the maximum mean signal in each dataset. The following datasets were used: 1805 (>Vg eye discs); 1800 (>H3.3K27M eye discs); and 1806 (>H3.3K27M >Vg eye discs).

#### For Figure 6A

Bedgraph files for H3K4me2 profiling were used for heatmapping around EPDnew promoters by the ‘computematrix’ with 10 bp binning and ‘plotHeatmap’ functions in deepTools. This promoter list is provided in Supplementary Information. Promoters were ordered by decreasing signal and displayed in the ‘YlGnBu’ color map. Datasets 1813 (w eye discs) and 1808 (w wing discs) were used.

#### For Figure 6B,C

UCSC Genome Browser tracks for H3K27me3 (red) and H3K4me2 (blue) datasets 1534 (w eye discs, H3K27me3), 1813 (w eye discs, H3K4me2), 1732 (w wing discs, H3K27me3), and 1808 (w wing discs, H3K4me2) were plotted with autoscaling.

#### For Figure 6D

Mean H3K4me2 signal across all EPDnew annotated promoters was rank-ordered for each sample and heatmapped using three-color conditional formatting in MS Excel with Red-Yellow-Green for low-to-high values. Selected values are shown in the figure. The full list is provided in Supplementary Information. Datasets 1813 (w eye discs); and 1808 (w wing discs) were used.

#### For Supplementary Figure 1A

Spearman correlations for all H3K27me3 profiling. Bedgraph files were binned into 5 kb windows, and correlations between the datasets were generated and displayed by the ‘plotcorrelation’ function in deepTools.

#### For Supplementary Figure 1B

Spearman correlations for all H3K4me2 profiling. Bedgraph files were binned into 500 bp windows, and correlations between the datasets were generated and displayed by the ‘plotcorrelation’ function in deepTools.

#### For Supplementary Figure 1C

Spearman correlations for all K27M profiling. Bedgraph files were binned into 5 kb windows, and correlations between the datasets were generated and displayed by the ‘plotcorrelation’ function in deepTools.

#### For Supplementary Figure 1D

Bedgraph files for K27M profiling were used for heatmapping around EPDnew promoters by the ‘computematrix’ with 10 bp binning and ‘plotHeatmap’ functions in deepTools. This promoter list is provided in Supplementary Information, Sheet 2. Promoters were ordered by decreasing signal and displayed in the ‘greys’ color map. 1802 (>H3.3K27M eye discs); and 1769 (>H3.3K27M >Vg eye discs) datasets were used.

#### For Supplementary Figure 1E

H3K4me2 signal was binned into 500 bp and displayed using the ‘multibigwigSummary’ and ‘plotPCA’ functions in deepTools.

#### For Supplementary Figure 1F

H3K27me3 signal was binned into 5 kb windows and displayed using the ‘multibigwigSummary’ and ‘plotPCA’ functions in deepTools.

**Supplementary Figure 1.**
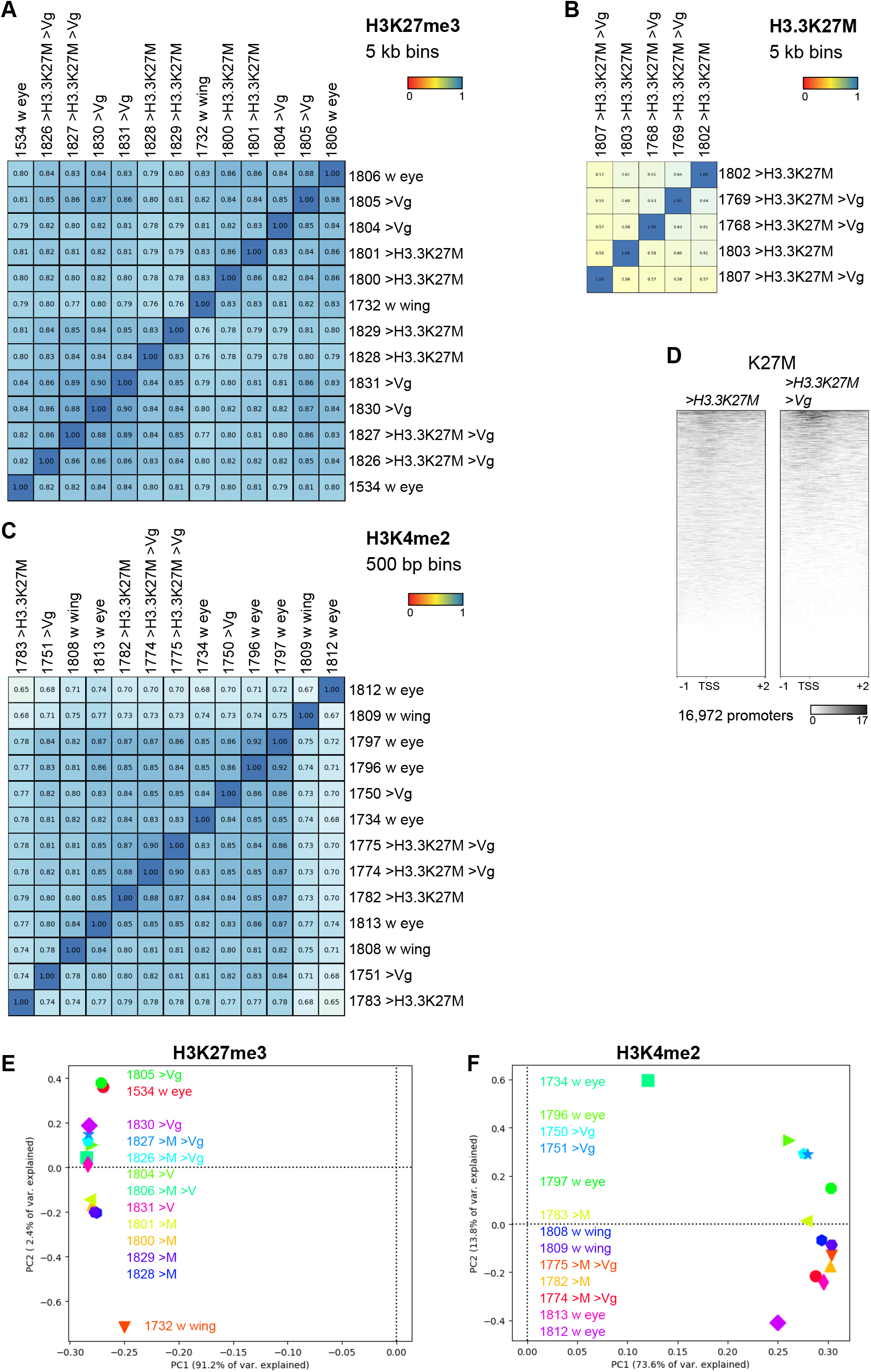
(**A**) Spearman correlations between H3K27me3 profiling samples. (**B**) Spearman correlations between H3K4me2 profiling samples. (**C**) Spearman correlations between K27M profiling samples. (**D**) Heatmap of K27M signal centered on promoters in >*H3.3K27M* and >*H3.3K27M* >*Vg* eye discs. (**E**) PCA plot of H3K4me2 signal in all samples. (**F**) PCA plot of H3K27me3 signal in all samples.

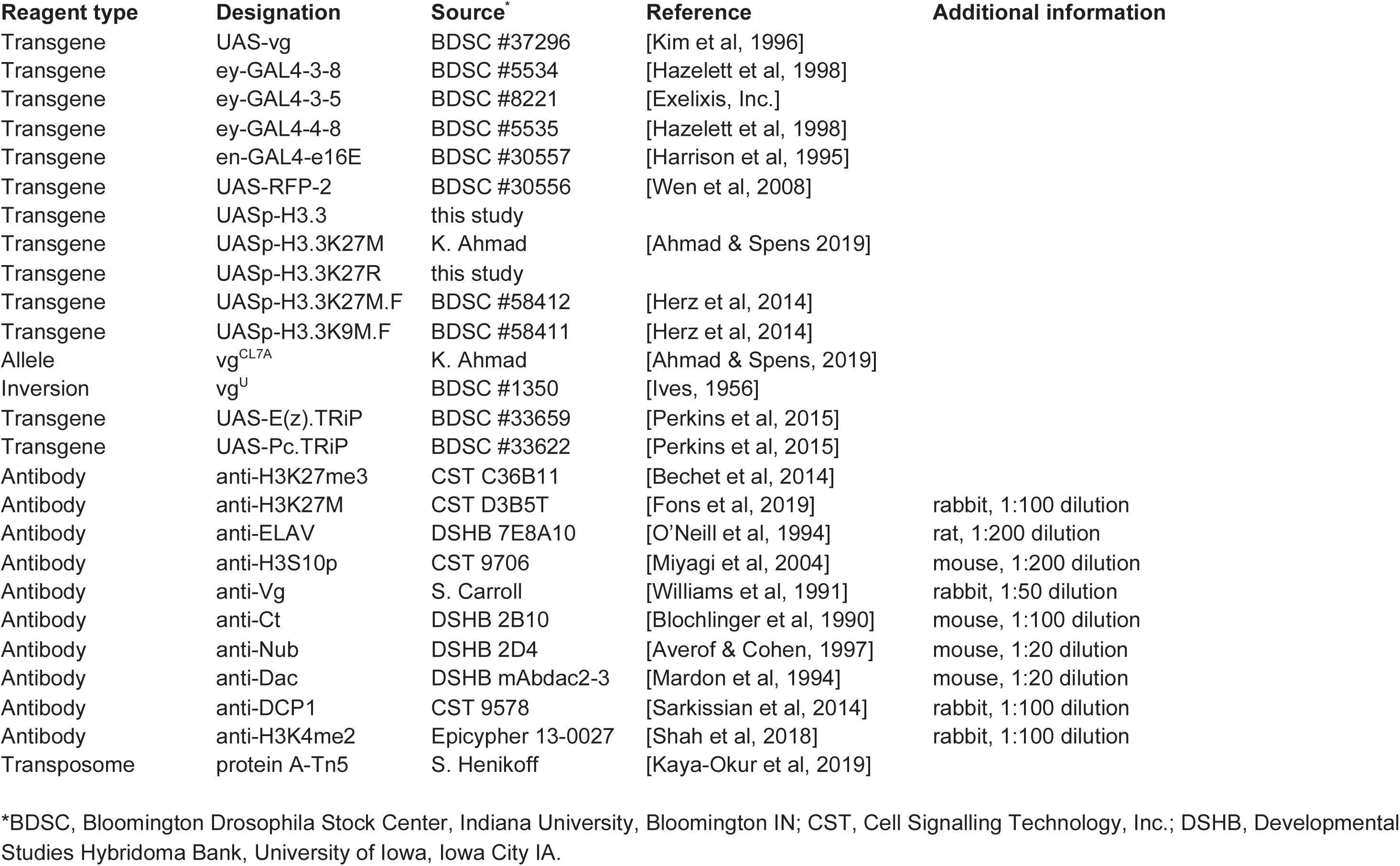
Key Resources Table.

